# Engineering low pH-tolerant *Komagataella phaffii* for industrial-ready itaconic acid production via methanol biorefinery

**DOI:** 10.1101/2025.01.29.634982

**Authors:** Fangqing Wei, Xiao Han, Fei Tao, Ping Xu

**Affiliations:** State Key Laboratory of Microbial Metabolism & School of Life Sciences & Biotechnology, Shanghai Jiao Tong University, Shanghai 200240, People’s Republic of China

**Keywords:** Methanol biorefinery, *Komagataella phaffii*, Itaconic acid, Synthetic biology, Low pH fermentation

## Abstract

Methanol biorefinery stands as a cornerstone strategy in the pursuit of sustainable biochemical production from non-food feedstocks. Despite its vast potential, bioconverting methanol into high titer biochemical production remains a formidable challenge. In this study, we have successfully engineered *Komagataella phaffii* to efficiently produce itaconic acid (IA), one of the top 12 building block chemicals, utilizing methanol as the sole carbon source. Our approach involved strategic metabolic engineering interventions—such as enhancing the supply of *cis*-aconitate and citrate precursors, minimizing isocitrate diversion, facilitating efficient formaldehyde assimilation, and optimizing IA export mechanisms—culminating in an unprecedented IA titer of 103.8 g/L in a 5-L bioreactor through fed-batch fermentation at pH 5.5. Remarkably, even at a reduced pH of 3.5, the strain still yielded 50.4 g/L of IA, highlighting the industrial viability of methanol biorefinery. This systematic work lays a robust foundation for high-titer production via methanol biorefinery, offering valuable insights for future research and applications.

## Introduction

Methanol has emerged as a pivotal renewable feedstock in the chemical manufacturing, offering a promising pathway towards sustainable and carbon-neutral society. Its liquid form ensures compatibility with existing infrastructure for transportation and fermentation, while its production from abundant sources like CO_2_ through photocatalytic or electrical reduction methods underscores its potential in establishing eco-friendly process chains. As the global energy demand surges, exacerbating environmental challenges, methanol stands out as an ideal C1 resource, capable of transforming into high-value chemicals through biotransformation, thereby addressing the urgent need for sustainable alternatives. Methanol possesses a higher degree of reducibility (with each carbon atom providing 50% more reducibility than glucose), implying that methanol can supply more electrons for the synthesis of target products^1,2^. Commercial facilities for green methanol production are currently in operation, with an anticipated annual output capacity escalating to approximately 8 million tons by 2027^3^. Notably, methanol presents a cost-effective alternative for other feedstocks of biorefinery. It is estimated that the price of green methanol will decrease to a range of 250–630 per ton by2050^4^. In contrast, glucose is priced around $540 per ton, and given its contention with food production sectors, this cost is foreseen to persistently increase^2,5,6^. Thus, the pursuit of non-food feedstocks like methanol has become crucial for the sustainable and economically viable synthesis of biochemicals^7^. Nonetheless, a prevailing challenge lies in the generally low titers of biochemical production achieved through methanol biorefinery processes (Table 1).

**Table 1.**
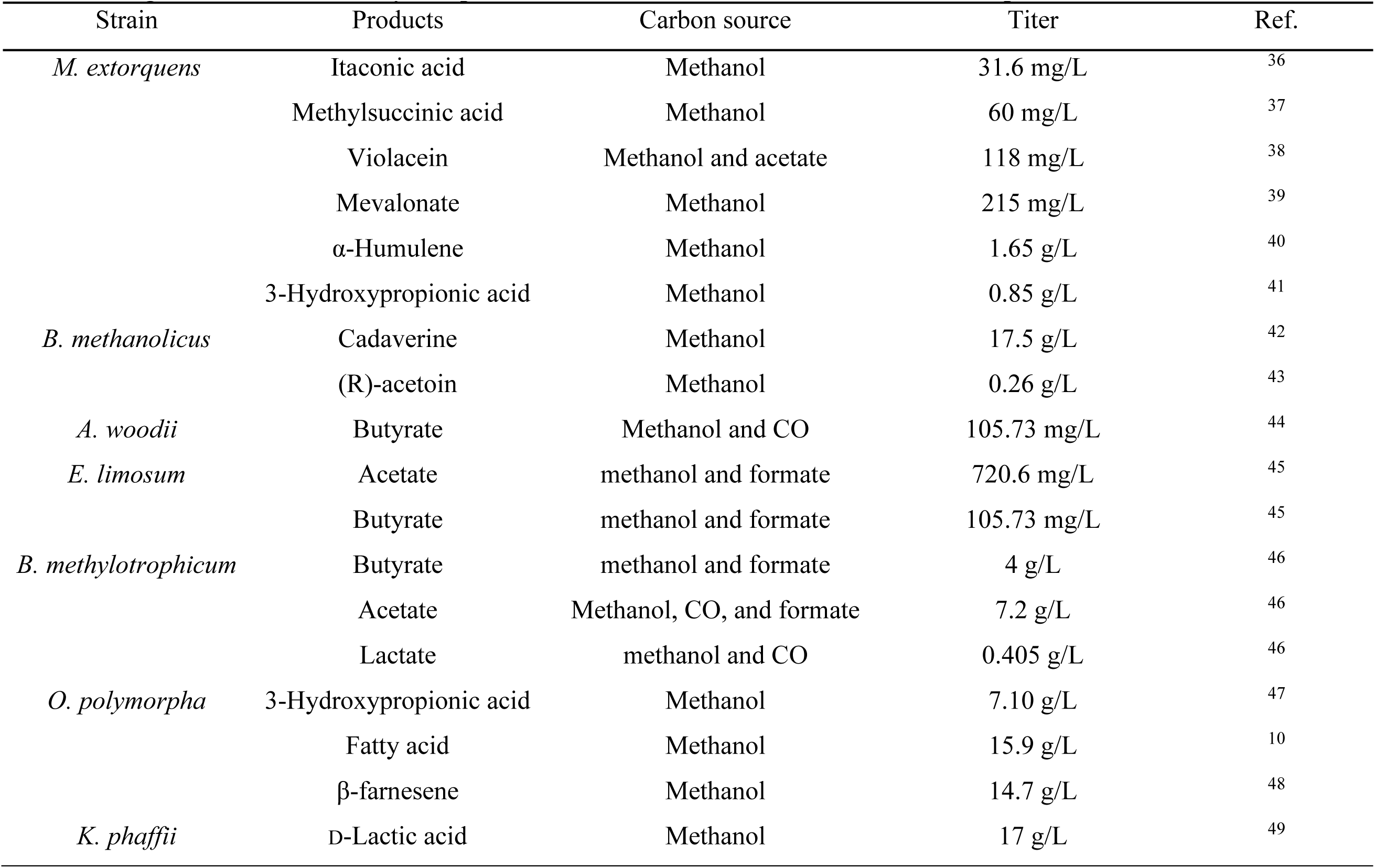
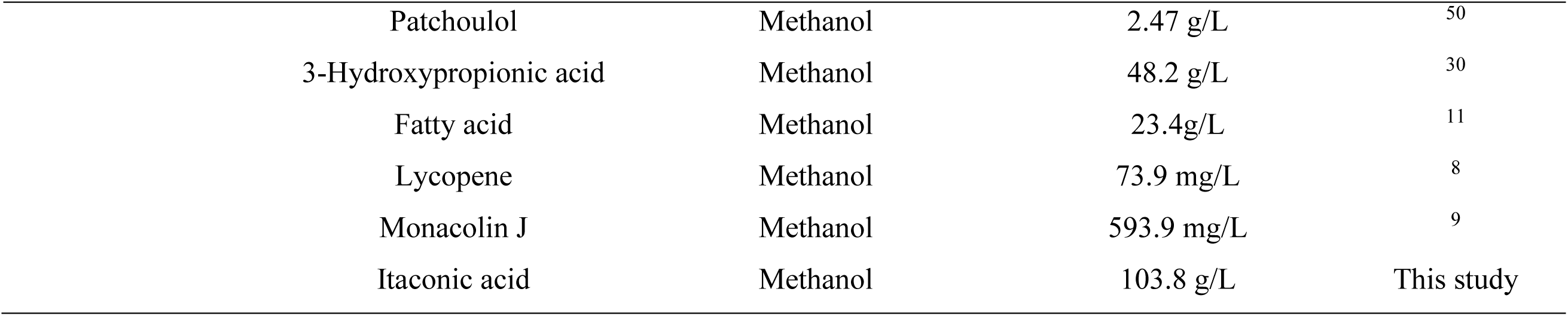
Engineered natural methylotrophs for methanol biotransformation to chemical products.

*Komagataella phaffii* (*K. phaffii*), recognized as a Generally Recognized as Safe (GRAS) organism, has been successfully engineered to synthesize an array of biochemicals, including lycopene^8^, monacolin J^9^, fatty alcohols^10^, and fatty acids^11^, utilizing methanol as the feedstock. The compartmentalized structures within *K. phaffii* cells, such as peroxisomes, plays a pivotal role in mitigating the cytotoxic effects of formaldehyde, thereby enhancing its robustness during metabolic processes. *K. phaffii* further distinguishes itself with its capability for high-density fermentation in cost-effective, minimal media, resilience against acidic environments, and inherent resistance to bacteriophage infections. These attributes collectively establish *K. phaffii* as a prime candidate for advancements in methanol biorefinery underscoring its potential in sustainable bioprocesses^12^.

Itaconic acid (IA), a C5 dicarboxylic acid derived from the TCA cycle, has been recognized by the U.S. Department of Energy as one of the top 12 most promising biobased platform chemicals^13^. IA is widely used in the synthesis of shape-memory polymers, adhesives, and antibacterial materials^14,15^. Recently, advancements have been made with the development of a biodegradable biobased itaconate ester rubber, further broadening the application spectrum of IA^16^. Currently, the production of IA by engineered organisms such as *Escherichia coli*, *Yarrowia lipolytica*, *Saccharomyces cerevisiae* and *Corynebacterium glutamicum* has been extensively studied^17–20^. However, these studies predominantly utilize glucose as a carbon source, which poses challenges due to its high cost and competition with food resources and land use which impedes the rapid progression of industrial biomanufacturing. Given that IA is a bulk chemical requiring substantial feedstock, it is more strategically sound to employ non-food sources like methanol, thereby mitigating competition with food production and promoting sustainability in industrial processes.

In this study, we systematically engineered *K. phaffii* for IA production from C1 feedstock, achieving unprecedented titers of 103.8 g/L at pH 5.5 and 50.4 g/L at pH 3.5 using methanol as the sole carbon source. This provides insights for the industrial scale-up of methanol biorefinery.

## Results

### Establishing the IA synthesis pathway in *K. phaffii*

In the natural IA-producing strain *Aspergillus terreus* (*A. terreus*), cytoplasmic *cis*-aconitate decarboxylase (encoded by the *At_cadA* gene) catalyzes the conversion of *cis*-aconitate to IA. This is the first reaction in the biosynthesis pathway of IA (Fig. 2a). Heterologous expression of *cadA* has successfully enabled non-native hosts such as *E. coli*^19^, *Y. lipolytica*^21^, and *S. cerevisiae*^17^ to produce IA. To enable IA production from methanol in *K. phaffii*, the *cis*-aconitate decarboxylase gene *At_cadA* was codon-optimized and expressed under the control of the strong methanol-inducible promoter *P_AOX1_* (alcohol oxidase 1 promoter). This expression cassette was integrated into the *PNSII-5* neutral integration site of the parental strain PC111^22^, resulting in the engineered strain PC-CA01.

**Fig. 1.**
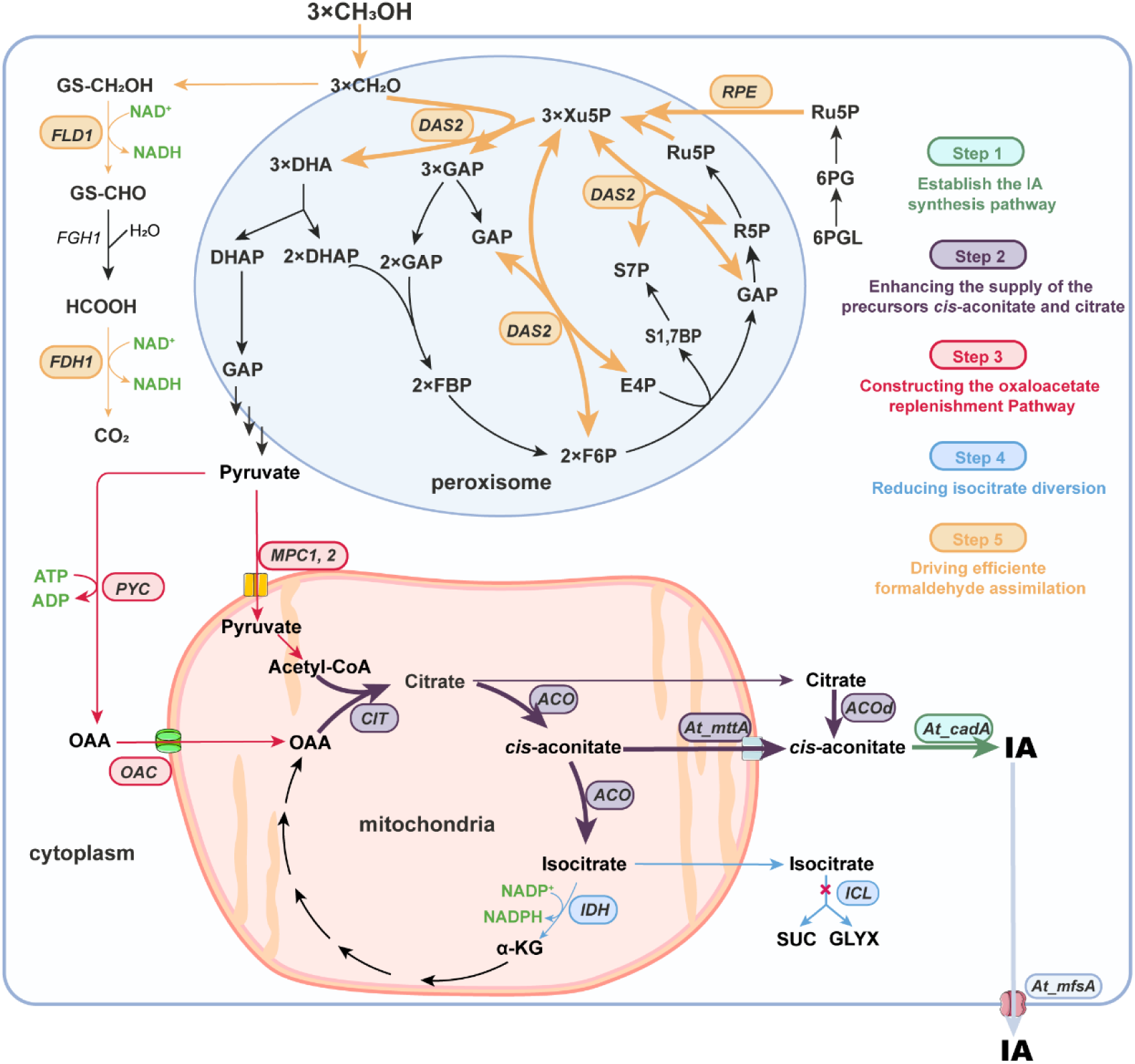
Metabolic engineering of *K. phaffii* by rewiring carbon metabolism to increase IA production. Five steps were designed to increase IA production: Step 1 (Establishing the IA synthesis pathway), Step 2 (Enhancing the supply of the precursors *cis*-aconitate and citrate), Step 3 (Constructing oxaloacetate replenishment pathway), Step 4 (Reducing isocitrate diversion), and Step 5 (Driving efficient formaldehyde assimilation). **Genes** that were overexpressed or modified are indicated in black italics. *At_cadA*: *cis*-aconitate decarboxylase gene from *A. terreus*, integrated into the *PNSII*-*5* locus. *At_mttA*: mitochondrial tricarboxylate transporter gene from *A. terreus*, integrated into the *PNSI*-*7* locus. *ACO*: aconitase gene, integrated into the *PNSI*-*9* locus, with *ACOd* representing *ACO* lacking the mitochondrial targeting sequence. *CIT*: citrate synthase gene, integrated into the *PNSI*-*8* locus. *IDH*: isocitrate dehydrogenase gene. *AMPD*: adenosine monophosphate (AMP) deaminase gene. *ICL*: isocitrate lyase gene. *At_mfsA*: major facilitator superfamily (MFS) transporter gene from *A. terreus*, integrated into the *PNSI*-*9* locus. *PYC*: pyruvate carboxylase gene, integrated into the *PNSIII*-*10* locus. *OAC*: oxaloacetate transporter gene, integrated into the *PNSIV*-*9* locus. *MPC*: mitochondrial pyruvate carrier proteins gene. *DAS2*: dihydroxyacetone synthase gene, integrated into the *PNSII*-*4* locus. *RPE*: endogenous D-ribulose 5-phosphate 3-epimerase gene, integrated into the *PNSII*-*9* locus. *FLD1*: formaldehyde dehydrogenase gene. *FDH1*: formate dehydrogenase gene. **Metabolites** are represented in black font. DHA: dihydroxyacetone. DHAP: dihydroxyacetone phosphate. GAP: glyceraldehyde-3-phosphate. Xu5P: xylulose 5-phosphate. Ru5P: ribulose-5-phosphate. R5P: ribose-5-phosphate. S7P: sedoheptulose-7-phosphate. S1,7BP: sedoheptulose-1,7-bisphosphate. E4P: erythrose-4-phosphate. FBP: fructose-1,6-bisphosphate. F6P: fructose-6-phosphate. 6PG: 6-phosphogluconate. 6PGL: 6-phosphogluconolactone. OAA: oxaloacetate. α-KG: α-ketoglutarate. SUC: succinic acid. GLYX: glyoxylic acid.

**Fig. 2.**
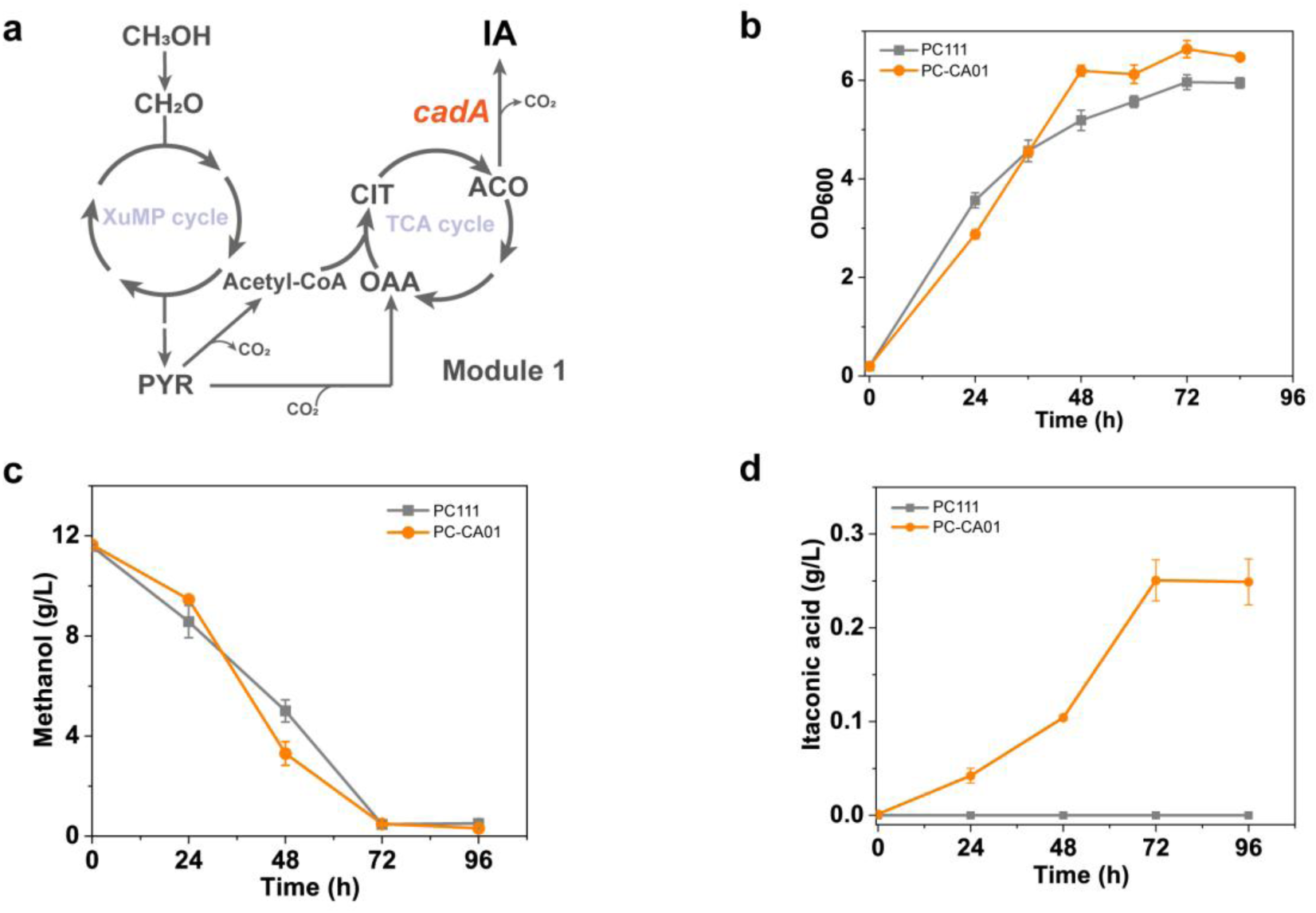
Establishing the IA synthesis pathway in *K. phaffii*. (a) Schematic representation of the metabolic pathway for IA synthesis from methanol. Overexpression of the codon-optimized *cis*-aconitate decarboxylase gene *cadA* from *Aspergillus terreus* resulted in the engineered strain PC-CA01. Modified genes are represented in orange-red font. Metabolites are represented in grey font. (b-d) Growth curves, methanol consumption, and IA production of the engineered strain PC-CA01 compared to the parental strain PC111 in shake flasks with methanol as the sole carbon source. All shake flask fermentations were conducted using Delft minimal medium with an initial methanol concentration of 10 g/L, at 30°C and 220 rpm for 72 h. Error bars correspond to the SD of the mean ± SD (n = 3, corresponding to three biological replicates).

In Delft minimal medium with methanol as the sole carbon source, the engineered strain PC-CA01 exhibited a growth pattern similar to the parental strain (Fig. 2b). After 72 hours of fermentation, all methanol was consumed, reaching an OD_600_ of 6.63 and producing 0.24 g/L of IA (Fig. 2b-d). In contrast, no IA was detected in the parental strain, indicating that the heterologous expression of *At_cadA* successfully enabled the production of IA from methanol. Notably, minimal amounts of common by-products such as ethanol, acetic acid, and formic acid were detected in the fermentation broth of both the engineered and parental strains.

### Enhancing the supply of the precursors *cis*-aconitate and citrate

*Cis*-aconitate, a precursor in the IA synthesis pathway, is primarily produced by the mitochondrial TCA cycle. However, the mitochondrial membrane structure significantly hinders the transport of *cis*-aconitate from the mitochondria to the cytosol, impeding IA synthesis. To address this issue, the endogenous mitochondrial tricarboxylate transporter from *A. terreus* (encoded by the *At_mttA* gene) was expressed to enhance the transmembrane transport of *cis*-aconitate between the mitochondria and the cytosol, resulting in the engineered strain MT02. MT02 showed a significant increase in IA production, reaching 1.21 g/L in shake flasks, which is 5 times higher than the control strain. Notably, the overexpression of *At_mttA* led to a 32% reduction in biomass (Fig. 3b), suggesting that it may have affected the mitochondrial membrane structure and, consequently, strain growth. Subsequently, the overexpression of endogenous citrate synthase gene *CIT* and aconitase gene *ACO* in *K. phaffii* was pursued to further enhance the synthesis of citrate and *cis*-aconitate. Overexpression of *CIT* (strain CT03) led to an increase in IA production, reaching 1.61 g/L, a 31.9% improvement over the control strain MT02 (Fig. 3c).

**Fig. 3.**
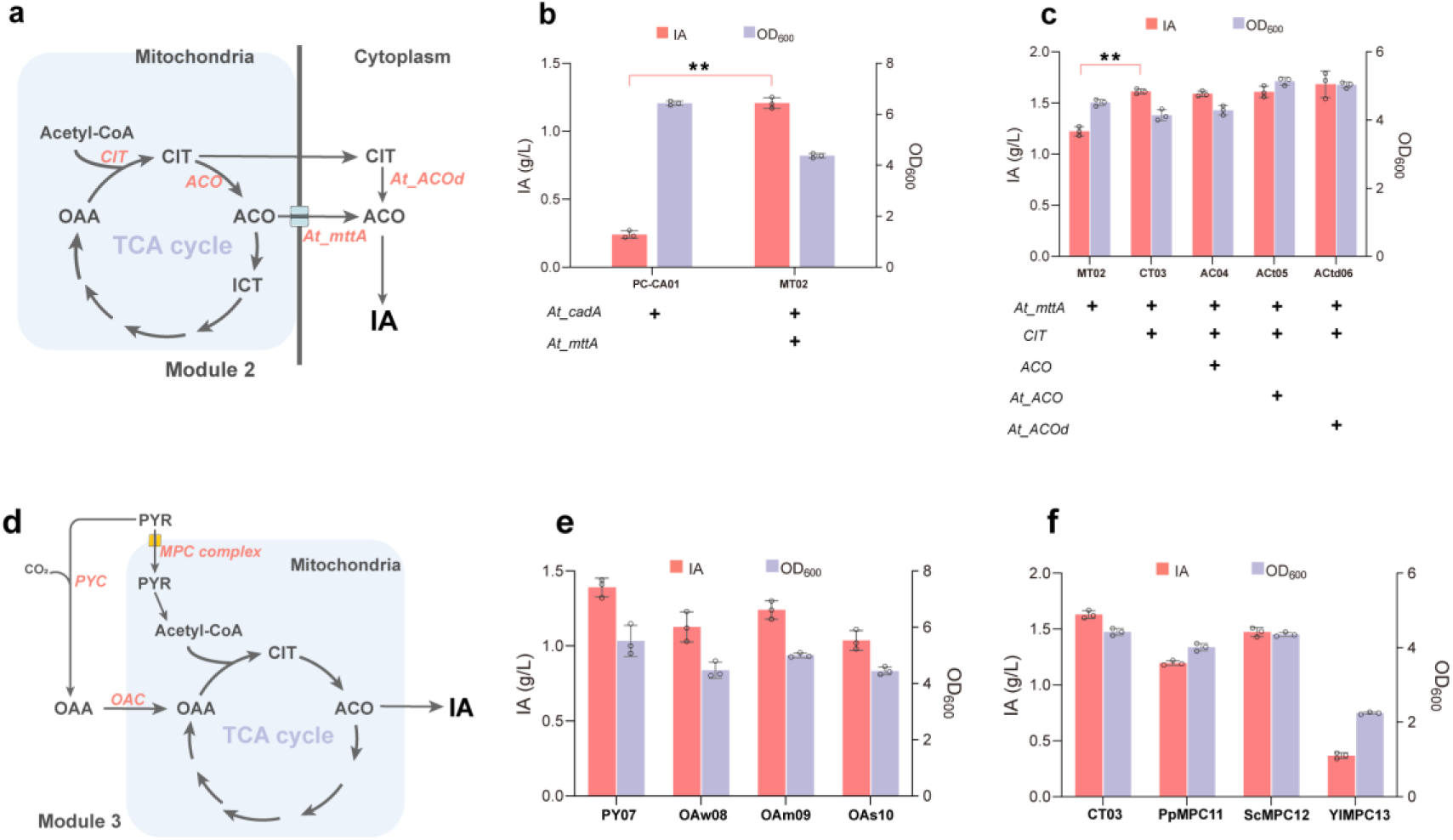
Enhancing precursor supply and constructing oxaloacetate replenishment pathway to increase IA production in *K. phaffii*. (a) Schematic diagram of metabolic engineering to enhance precursor supply. Modified genes are represented in orange-red font. Metabolites are represented in grey font. Overexpression of the mitochondrial tricarboxylate transporter gene *mttA* (strain MT02) and the aconitase gene *ACO* (strain Act05) from *Aspergillus terreus*, the endogenous citrate synthase gene *CIT* from *K. phaffii* (strain CT03), and the aconitase gene *ACO* (strain AC04) was performed. The ACtd06 strain represents the *ACO* sequence with the mitochondrial targeting sequence removed. (b) Titers of IA and cell density of strains after overexpressing *At_mttA.* (c) Titers of IA and cell density of strains after enhancing precursor supply. (d) Schematic diagram of metabolic engineering for constructing the oxaloacetate replenishment pathway. Overexpression of the endogenous pyruvate carboxylase gene *PYC* (strain PY07), mitochondrial pyruvate carrier proteins from different sources (strains PpMPC11, ScMPC12, YlMPC13), and the expression of the oxaloacetate carrier gene *OAC* driven by promoters of varying strengths was performed. (e) Titers of IA and cell density of strains after overexpressing *OAC*. (f) Titers of IA and cell density of strains after overexpressing *MPC*. Error bars correspond to the SD of the mean ± SD (n = 3, corresponding to three biological replicates). Statistical significance of the different IA levels in comparison with the control was evaluated using Student’s t-test (**\***, P < 0.05; **\*\***, P < 0.01).

However, adding an additional copy of *ACO* (strain AC04) did not change production levels, suggesting that the endogenous *ACO* activity might be weak. For this reason, aconitase sourced from *A. terreus* was heterologously expressed. Simultaneously, to fully utilize the cytosolic citrate, a cytosolic aconitase pathway was constructed (Fig. 3a). The aconitase gene from *A. terreus* was codon-optimized and its mitochondrial targeting sequence (MTS) was predicted using MitoFates (https://mitf.cbrc.pj.aist.go.jp/MitoFates/cgi-bin/top.cgi). Based on these predictions, the first 18 amino acids of the aconitase were removed (*At_ACOd*) to create strain ACtd06, while a strain retaining the MTS served as a control (strain ACt05). As shown in Fig. 3c, the construction of the cytosolic aconitase pathway slightly increased the IA production by 4.3%, reaching 1.68 g/L. However, the production remained almost unchanged with the mitochondrially localized aconitase, suggesting that the expression level of mitochondrial aconitase might already be sufficient to catalyze the conversion of citrate within the mitochondria effectively.

### Constructing the oxaloacetate replenishment pathway

The TCA cycle begins with the condensation of acetyl-CoA and oxaloacetate to form citrate, and the availability of the oxaloacetate could be a limiting factor in TCA cycle efficiency. To address this, we enhanced the synthesis of oxaloacetate by overexpressing the endogenous pyruvate carboxylase in *K. phaffii* (strain PY07). Additionally, we overexpressed the endogenous oxaloacetate carrier protein OAC in *K. phaffii*, identified through a BLAST search against the oxaloacetate carrier protein from *S. cerevisiae* (Sc_OAC), a mitochondrial inner membrane transporter. Deletion of OAC in *S. cerevisiae* has been reported to reduce mitochondrial oxaloacetate content^23^. However, the IA production in the engineered strain PY07 decreased by 13.6%, possibly due to the overexpression of pyruvate carboxylase, which diverted some pyruvate away from entering the mitochondria (Fig. 3e). Subsequently, we expressed OAC using weak, medium, and strong promoters (*P_CAT1_*, *P_DAS1_* and *P_PMP20_*), but the IA production in the engineered strains (OAw08, OAm09, OAs10) decreased by 18.7%, 10.7%, and 25.1%, respectively. Correspondingly, biomass decreased by 18.9%, 9.2%, and 19.4%. Oxaloacetate is closely linked with the NADH shuttle system, and its overexpression could potentially disrupt the balance of energy metabolism^24^.

Pyruvate generated during methanol assimilation predominantly resides in the cytosol and must be transported across the membrane into the mitochondria, where it is converted into acetyl-CoA by the pyruvate dehydrogenase complex, entering the TCA cycle. To facilitate increased transport of pyruvate into the mitochondrial TCA cycle, mitochondrial pyruvate carrier proteins (MPC1 and MPC2) were overexpressed^25^. Transporter proteins from *K. phaffii*, *S. cerevisiae*, and *Y. lipolytica* were selected to create the strains PpMPC11, ScMPC12, and YlMPC13, respectively. As shown in the Fig. 3f, the production of IA in these strains decreased by 26.3%, 9.8%, and 77.3%, respectively. Notably, strain YlMPC13 experienced significant growth inhibition, with a 49.3% reduction in biomass, indicating that overexpression of this transporter imposes a substantial burden on strain growth.

### Reducing isocitrate diversion

Reducing the isocitrate diversion helps accumulate the precursor *cis*-aconitate. Here, we downregulated isocitrate dehydrogenase (IDH) expression and knocked out the glyoxylate shunt. Two strategies were used to reduce IDH expression: first, replacing its native promoter with weaker ones (*P_TPI_*, *P_ADH2_*, *P_FBA1_*), which resulted in the strains IDp14, IDp15, and IDp16, respectively; second, overexpressing adenosine monophosphate (AMP) deaminase (AMPD) (strain IDp17), which converts AMP to inosine monophosphate (IMP), thereby inhibiting the AMP-dependent activity of IDH^26^. As shown in the Fig. 4b, replacing the native promoter with the weak promoters *P_TPI_* and *P_ADH2_* resulted in further increases in IA production by 12.2% and 4.9%, respectively, with the *P_TPI_* strain achieving 1.83 g/L in shake flasks. However, replacing the promoter with *P_FBA1_* led to a 6.1% decrease in IA production. Additionally, overexpression of endogenous AMPD did not enhance IA production.

**Fig. 4.**
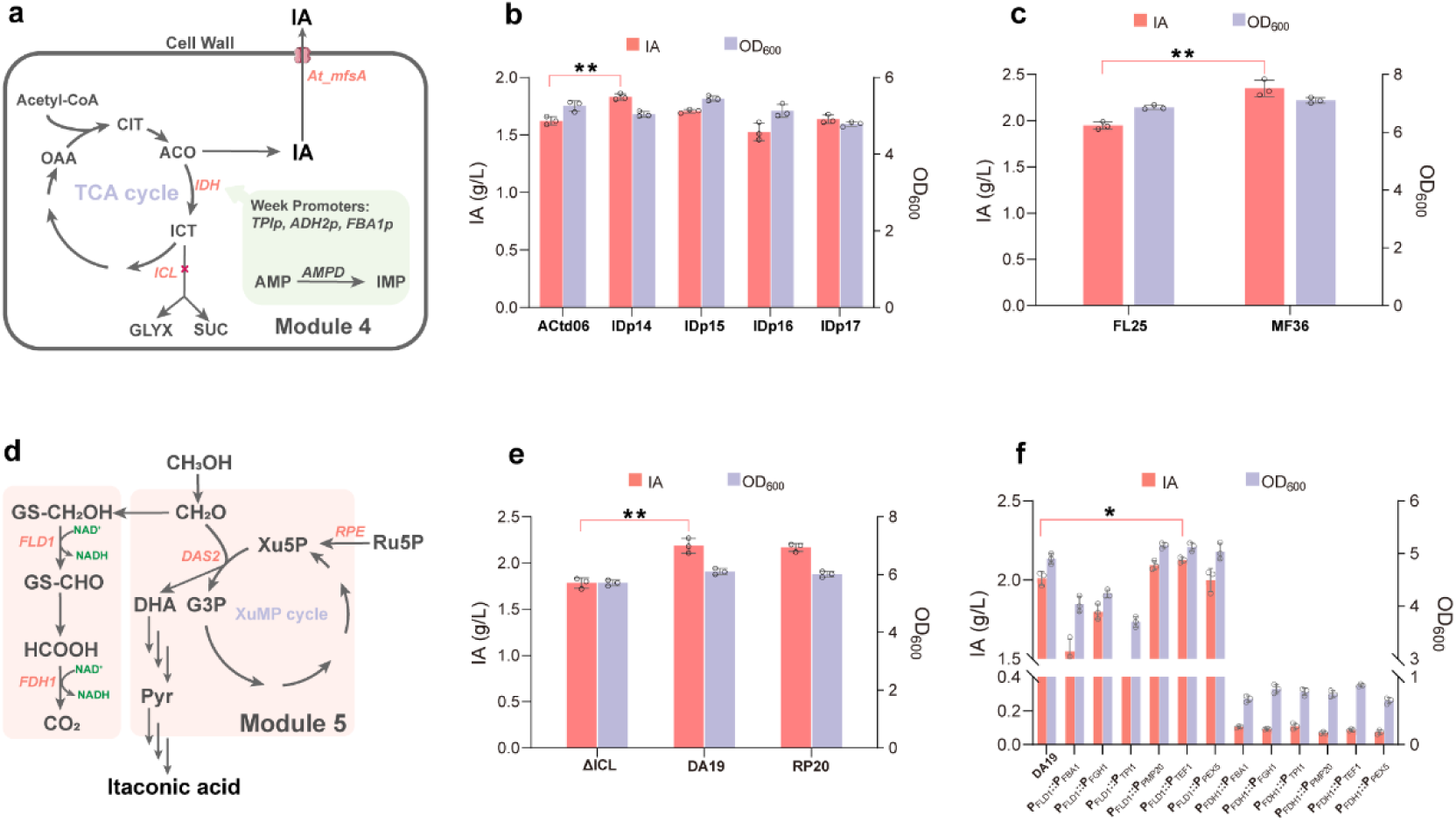
Reducing isocitrate diversion and driving efficient formaldehyde assimilation to increase IA production. (a) Schematic diagram of metabolic engineering for reducing isocitrate diversion. Overexpression of *AMPD* and the use of a weak promoter replacement strategy (strains IDp14, IDp15, IDp16, IDp17) were employed to downregulate the expression of isocitrate dehydrogenase (*IDH*), along with the knockout of the isocitrate lyase gene (*ICL*). Modified genes are represented in orange-red font. Metabolites are represented in grey font. (b) Titers of IA and cell density of strains after downregulating the *IDH* expression. (c) Titers of IA and cell density of strains after overexpressing *At_mfsA*. (d) Schematic diagram of metabolic engineering for driving efficient formaldehyde assimilation. Overexpression of *DAS2* (strain DA19) and *RPE* (strain RP20) was conducted, along with the downregulation of *FLD1* and *FDH1* expression using weak promoters. (e) Titers of IA and cell density of strains after overexpressing *DAS2* and *RPE*. (f) Titers of IA and cell density of strains after downregulating *FLD1* and *FDH1*. Error bars correspond to the SD of the mean ± SD (n = 3, corresponding to three biological replicates). Statistical significance of the different IA levels in comparison with the control was evaluated using Student’s t-test (**\***, P < 0.05; **\*\***, P < 0.01).

A portion of isocitrate in the mitochondria crosses the membrane into the cytosol, where it is cleaved by isocitrate lyase (ICL) into glyoxylate and succinate, subsequently entering the glyoxylate cycle^27^. To reduce carbon flux through the glyoxylate shunt, we knocked out ICL. Using the isocitrate lyase gene *ICL1* from *S. cerevisiae* as a template, we performed a BLAST search to identify the endogenous ICL gene in *K. phaffi* and constructed a ΔICL strain. After the knockout, the biomass of the strain slightly decreased, but the IA production did not increase (Fig. S1).

### Driving efficient formaldehyde assimilation

After methanol is oxidized to formaldehyde, in addition to being assimilated into the central carbon metabolism, a portion is further oxidized via the dissimilation pathway to CO_2_, providing energy and detoxification for the cells. However, the presence of the dissimilation pathway results in more than 50% of the carbon atoms being wasted as CO_2_, leading to low conversion yield in methanol bioconversion^7,28^. Therefore, it is necessary to enhance methanol assimilation to improve the product conversion yield. The dihydroxyacetone synthase gene *DAS2* catalyzes the condensation of formaldehyde and xylulose-5-phosphate (Xu5P) to form dihydroxyacetone. Overexpressing this gene in the engineered strain DA19 resulted in a 7.1% increase in biomass and a 22.4% increase in IA production, reaching 2.18 g/L in shake flasks (Fig. 4e). Next, we attempted to increase the supply of Xu5P by overexpressing *RPE* (strain RP20), which encodes D-ribulose-5-phosphate 3-epimerase. However, this did not lead to a significant change in IA production (Fig. 4e).

Next, we aim to weaken the dissimilatory pathway to further improve IA yield. Given that knocking out *FLD1* or *FDH1* severely impacts cell growth^28,29^, we adopted the strategy of replacing their native promoters with weaker promoters, as described by Wu et al.^30^. We replaced the *FLD1* and *FDH1* promoters with promoters of varying strengths: 10% (*P_FBA1_*, *P_FGH1_*), 30% (*P_TPI1_*, *P_PMP20_*), and 60% (*P_TEF1_*, *P_PEX5_*) of the original *P_FLD1_* strength. As shown in the Fig. 4f, replacing *P_FLD1_* with *P_FBA1_*, *P_FGH1_*, and *P_TPI1_* resulted in production decreases of 22.8%, 10.4%, and 43.2%, respectively. In contrast, replacing *Pfld1* with *Ppmp20* and *Ptef1* increased production by 4.0% and 5.9%, respectively, while *Ppex5* showed no significant change in production. When *Pfdh1* was replaced, all strains exhibited severely impaired growth on methanol, with biomass after 72 h of fermentation ranging from 0.7 to 0.9, indicating that *FDH1* is highly sensitive for methanol metabolism.

Transcription factors can be used to regulate the expression levels of genes. In designing the metabolic pathway for IA production from methanol, several nodes (*AOX1*, *CIT*, and *At_cadA*) utilize the methanol-inducible strong promoter *P_AOX1_* (Fig. S2a). By expressing transcriptional activators of this promoter, it is anticipated that the expression levels of these nodes can be globally enhanced. Preliminary research has been conducted on the transcriptional regulation system of the *P_AOX1_*, identifying transcription activators such as *Mit1*, *Mxr1*, and *Prm1*^31,32^. However, overexpression of the transcription activators *Mit1* (strain Mt33) and *Mxr1* (strain Mr34) did not increase IA production. In contrast, overexpression of *Prm1* (strain Pm35) resulted in a 29.2% decrease in biomass and a 30.1% reduction in titer (Fig. S2b).

### Fed-batch fermentation for IA production

To evaluate the potential of the engineered strain for IA production using methanol, fed-batch fermentation was conducted in a 5-L bioreactor. Fermenting with methanol as the sole carbon source for 144 h in the 5-L bioreactor, the biomass (OD_600_) reached 124, and 62.4 g/L of IA was produced with a conversion yield of 0.26 g/g, representing 38.4% of the theoretical yield (Fig. 5b). This production level is 26.9 times higher than that achieved in shake flasks.

**Fig. 5.**
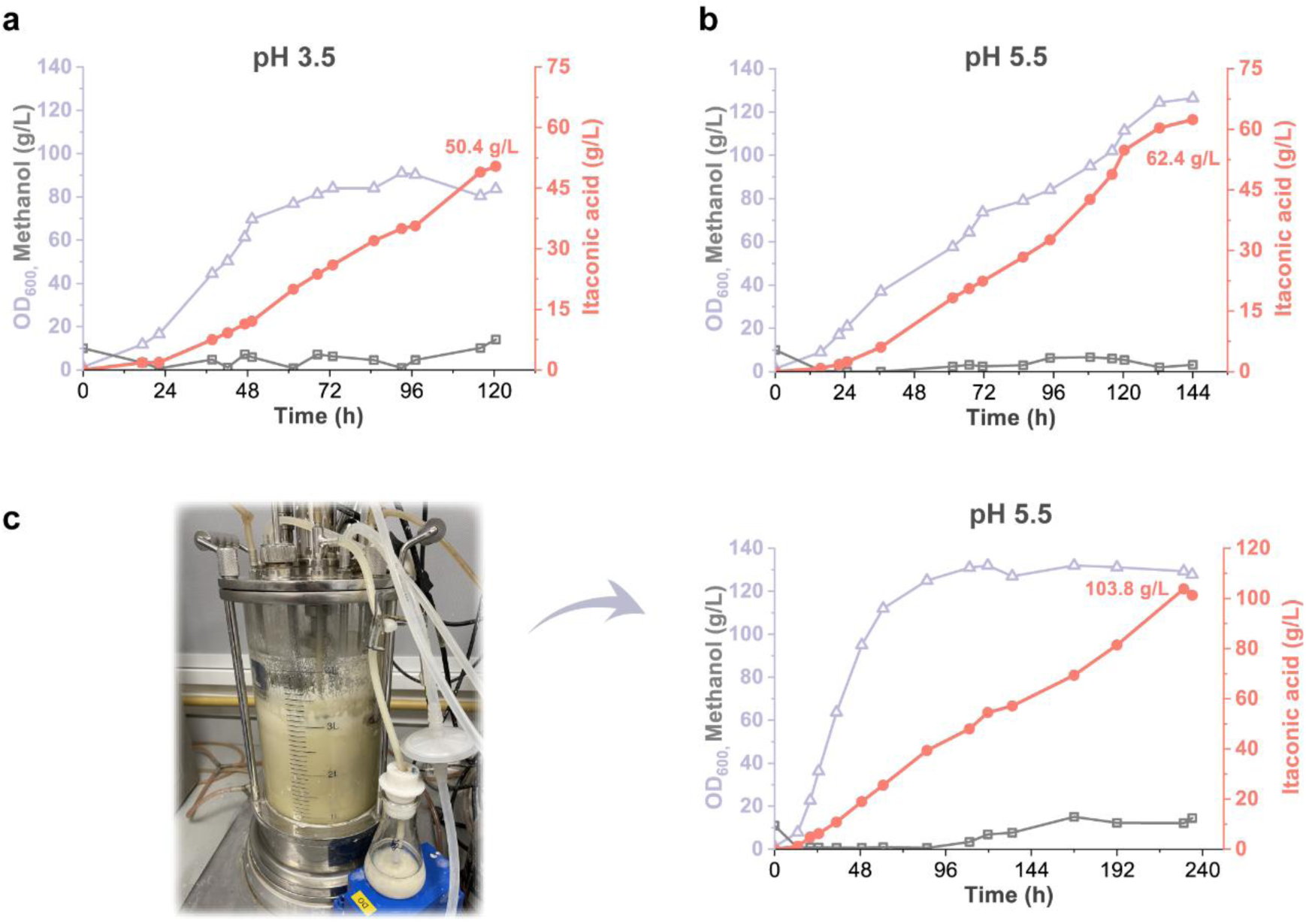
Fed-batch fermentation for IA production. (a) Low pH fed-batch fermentation. KOH was used to control the pH at 5.5 for the initial 36 h to accumulate biomass, after which the fermentation was allowed to naturally adjust to and maintain pH 3.5. (b) Fed-batch fermentation with pH controlled at 5.5 using KOH. (c) Fed-batch fermentation with pH controlled at 5.5. Batch fermentation was conducted in a 5-L bioreactor at 30°C, with dissolved oxygen levels maintained above 25%.

Low pH fermentation can reduce the cost of neutralizing agents. Additionally, when the pH is below the pKa values of IA’s two carboxyl groups (3.85 and 5.45), IA primarily exists in its non-dissociated form, simplifying downstream separation processes. Consequently, we conducted fed-batch fermentation at pH 3.5. To shorten the fermentation period, KOH was used to control the pH at 5.5 for the initial 36 h to accumulate biomass, after which the fermentation was allowed to naturally adjust to and maintain pH 3.5. As shown in Fig. 5a, after 72 h of fermentation, the biomass entered a stable phase, reaching an OD_600_ of 84, which represents a 32.2% reduction compared to fed-batch fermentation at pH 5.5. After 120 h of fermentation, the maximum IA titer reached 50.4 g/L, only a 19.2% reduction compared to the pH 5.5 fermentation. The yield of IA was 0.24 g/g, demonstrating that the IA production capability of *K. phaffii* is only minimally affected under low pH conditions, confirming its broad applicability as a host for organic acid production.

To further increase the titer of IA, we optimized fermentation conditions. As shown in Fig. 5c, after 60 h of fermentation, the biomass stabilized at an OD_600_ of 112. At 230 h, the IA titer peaked at 103.8 g/L, with a biomass of 129.3, yielding 0.23 g/g. To our knowledge, this is the highest titer of a biochemical produced from methanol reported in the literature.

## Discussion

Using non-food feedstocks for biochemical production can help avoid two major issues associated with food feedstocks: First, food feedstocks are costly, often accounting for over 50% of total production costs^5^. Second, using food as a feedstock raises the issue of competition for food resources. As the demand for feedstocks in biochemical production, especially for bulk chemicals, continues to grow, the development of non-food alternatives has become essential. Methanol is an ideal non-food feedstock, which can be synthesized from CO_2_, with abundant reserves, and its costs are expected to decrease significantly (prices of green methanol are projected to drop to between $250 and $630 per ton by 2050)^4^. However, the titer of biochemicals produced through methanol biorefinery is generally low, falling significantly short of the requirements for industrial production (Table 1). In this study, we employed rational metabolic engineering to design *K. phaffii* for the efficient production of IA using methanol as the sole carbon source, achieving a titer of 103.8 g/L, which is close to the industrial production levels obtained with glucose. These results demonstrate the industrial potential of methanol biorefinery for producing bulk chemicals and hold promise for replacing food feedstocks like glucose.

Rational design in metabolic engineering has led to a gradual increase in IA production. Three factors may contribute to achieving high titers of IA: (a) Reducing formaldehyde accumulation. Overexpression of the dihydroxyacetone synthase gene (*DAS2*) reduces formaldehyde buildup, enhancing both cell growth and product yield, with similar outcomes observed in studies on fatty acid production^10,11^. (b) Enhancing the supply of precursors, *cis*-aconitate, and citrate. Overexpression of citrate synthase, aconitase, the mitochondrial tricarboxylate transporter, and reduction of isocitrate diversion enhance the supply of precursors, significantly boosting production. (c) Strengthening transmembrane transport processes. The synthesis of IA in *K. phaffii* involves two transmembrane transport processes: transporting *cis*-aconitate from the mitochondria to the cytosol where it is decarboxylated to form IA, and transporting IA from the cytosol to the extracellular space. These two processes were enhanced by overexpressing the transporter *mttA* and *mfsA*.

*K. phaffii*, capable of tolerating low pH and resisting phage infections^12^, holds significant potential as a host for organic acid production. However, there is currently a lack of research on organic acid fermentation at low pH using methanol as the substrate. We conducted fed-batch fermentation of *K. phaffi* at pH 3.5. Under these conditions, the need for neutralizing agents was significantly reduced, and IA remained primarily in its non-dissociated form, greatly simplifying downstream processing^34^. The IA titer reached 50.4 g/L (Fig. 7a), a decrease of only 19.2% compared to fed-batch fermentation at pH 5.5 (Fig. 7b). This indicates that *K. phaffi’*s IA production capacity is only slightly affected under low pH, demonstrating its versatility as a host for organic acid production. This is the first instance of low-pH fermentation using methanol as the sole carbon source, offering insights for methanol biorefinery that aim to produce other organic acids without neutralizing agents or at low pH.

In summary, by employing rational design in metabolic engineering of *K. phaffii*, we enhanced the supply of precursors such as citrate and *cis*-aconitate, weakened the isocitrate diversion, and drove efficient methanol assimilation. The engineered strain achieved a production titer of 103.8 g/L of IA in a 5 L bioreactor, approaching industrial level. Under pH 3.5 conditions, the low pH fermentation resulted in a production of 50.4 g/L of IA, demonstrating the significant potential of *K. phaffii* for organic acid production without neutralizing agents. These results provide insights for the production of bulk chemicals and other biochemicals through methanol biorefinery.

## Acknowledgements

This research was supported by the National Research Foundation, Prime Minister’s Office, Singapore under its Campus for Research Excellence and Technological Enterprise (CREATE) programme, and the National Natural Science Foundation of China (Grant No. 32170105). The authors also thank the Double First Class Plan of Shanghai Jiao Tong University for the financial support provided.

The authors would like to thank Prof. Yongjin J. Zhou and PhD. Peng Cai from the Dalian Institute of Chemical Physics for providing the starting strain PC111 and the plasmid pCAI-gPNSI-1.

## Author contributions

Fei Tao, Fangqing Wei, Xiao Han, and Ping Xu designed the experiments. Fangqing Wei performed the experiments. Fangqing Wei wrote the manuscript. Xiao Han, Fei Tao and Ping Xu revised the manuscript. Ping Xu and Fei Tao conceived the projects.

---

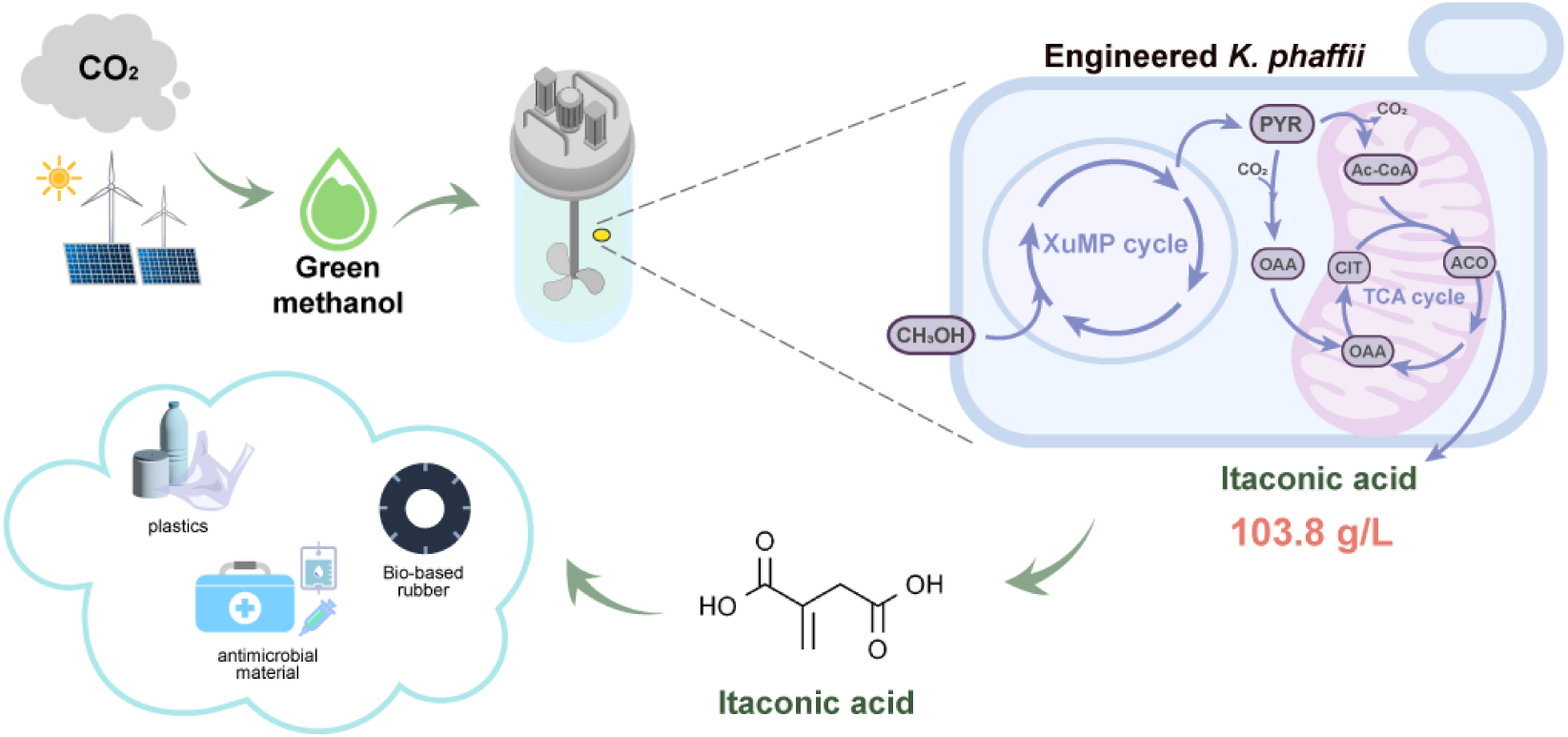
Schematic of itaconic acid production via methanol biorefinery. Methanol could be synthesized from CO_2_, which is subsequently converted into itaconic acid within microbial cell factories. This process serves as a model for non-food feedstock alternatives and contributes to carbon neutrality efforts.

## Materials and methods

### Strains, plasmids, and media

All strains and plasmids used in this study are listed in Tables S1-3. Sequences of genes, promoters and terminators used in this study are listed in Tables S4. *E. coli* DH5α, employed for gRNA plasmid construction, was cultivated in LB medium (5 g/L yeast extract, 10 g/L tryptone, 10 g/L NaCl) at 37 °C with shaking at 220 rpm. The engineered *K. phaffii* strains were developed from the *HIS4* auxotrophic *K. phaffii* PC111^22^, which was kindly provided by Prof. Yongjin J. Zhou from the Dalian Institute of Chemical Physics, Chinese Academy of Sciences. Other engineered *K. phaffii* strains were constructed in this study. Delft minimal medium (2.5 g/L (NH_4_)_2_SO_4_, 14.4 g/L KH_2_PO_4_, 0.5 g/L MgSO_4_·7H_2_O, 20 g/L glycerol, or 10 g/L methanol, 40 mg/L histidine, 1 ml/L Vitamin solution, 2 ml/L Trace metal solution) and YPD medium (10 g/L yeast extract, 20 g/L tryptone and 20 g/L glucose) were used for cultivating *K. phaffii*, which were incubated at 30 °C with shaking at 220 rpm. During fed-batch fermentation in a 5-L bioreactor, 5 g/L of yeast extract was used instead of vitamin solutions and trace metal solutions. Methanol and itaconic acid were purchased from Aladdin (Shanghai). DNA oligonucleotide primers were synthesized by Tsingke Biotech (Shanghai, China).

### DNA manipulation

Gene insertion and deletion were achieved through co-transformation of gRNA plasmids and donor fragments. The 20 bp target sequences in the gRNA plasmids were designed using the web toolbox (http://chopchop.cbu.uib.no). For instance, in the knockout of the isocitrate lyase gene (*ICL*), primers gICL-F and gICL-R were used to obtain the plasmid pCAI-gICL by PCR, using the pCAI-gRNA plasmid as a template. The PCR product was transformed into *E. coli* DH5α competent cells and plated on LB agar containing 50 ng/µL kanamycin for selection of transformants and plasmid extraction. The donor fragments were constructed using overlap extension PCR. Initially, the upstream homologous arm ICL-up and the downstream homologous arm ICL-dn were amplified from the *K. phaffii* GS115 genome using primers ICL up-F/ICL up-R and ICL dn-F/ICL dn-R, respectively. Subsequently, the two homologous arms were fused using primers ICL up-F/ICL dn-R to produce the ICL donor fragment. All endogenous gene fragments, promoters, and terminators were amplified by PCR using genomic DNA from *K. phaffii* GS115 as the template. Exogenous gene fragments and terminators were obtained by PCR amplification using genomic DNA from *S. cerevisiae* S288C and *Y. lipolytica*. Genes *cadA*, *mttA*, and *mfsA*, originating from *A. terreus*, were codon-optimized and synthesized by Tsingke Biotech (Shanghai, China).

### Batch and Fed-Batch Fermentation

Shake flask batch fermentations for production of itaconic acid were carried out in a 100 mL shaking flask with 20 mL Delft minimal medium. The genetically engineered strains were initially cultured overnight in YPD medium. Subsequently, they were transferred to Delft minimal medium supplemented with 20 g/L glycerol and 10 g/L methanol and cultured for an additional 24 hours. After this preparatory phase, the cultures were inoculated into shake flasks using methanol as a carbon source for fermentation, maintaining a final optical density (OD_600_) of 0.2. During shake flask fermentation, an appropriate amount of 4M KOH is added every 24 h to maintain the pH above 5.0.

The fed-batch fermentation in bioreactors was conducted in a 5-L fermentation bioreactor (Bailun Bio, Shanghai, China) containing 2.7 L of fresh Delft minimal medium, and the inoculation volume was 10 % (v/v), so that the final initial fermentation liquid volume was 3 L. The temperature, agitation, aeration, and pH were monitored and controlled using a BLBIO-B control system (Bailun Bio, Shanghai, China). The temperature was kept at 30 °C, and the initial agitation was set to 100 rpm and increased to maximally 800 rpm depending on the dissolved oxygen (DO) level^22^. The dissolved oxygen level was maintained above 25%, and the pH was kept at pH 5.5 by automatic addition of 4 M KOH. During the fed-batch cultivation, the cells were fed with 600 g/L methanol and 10 × supplementary components (25 g/L (NH_4_)_2_SO_4_, 144 g/L KH_2_PO_4_, 5 g/L MgSO_4_·7H_2_O, 10 × trace metal, and 10 × vitamin solutions)^11^. The feeding rate of methanol solution was adjusted from 40 g/L/d to120 g/L/d, keeping the methanol concentration below 20 g/L. The feeding of supplementary component solution was set as half of the feeding speed of methanol solution.

### Analytical methods

The cell density (OD_600_) was measured by a spectrophotometer. The concentrations of methanol, itaconic acid, formic acid, acetic acid and ethanol were measured via a high-performance liquid chromatography (HPLC) system equipped with a Bio-Rad Aminex HPX-87H column and a differential refractive index detector^11^. The samples were eluted with 5 mM H_2_SO_4_ at a flow rate of 0.5 mL/min at 55 °C for 30 min.

## Reference

1. Lais, A., Gondal, M.A., Dastageer, M.A. & Al-Adel, F.F. Experimental parameters affecting the photocatalytic reduction performance of CO_2_ to methanol: A review. Int. J. Energy Res. 42, 2031−2049 (2018).

2. Zhang, W. et al. Guidance for engineering of synthetic methylotrophy based on methanol metabolism in methylotrophy. RSC Adv. 7, 4083−4091 (2017).

3. Antoniewicz, M.R. Synthetic methylotrophy: Strategies to assimilate methanol for growth and chemicals production. Curr. Opin. Biotechnol. 59, 165−174 (2019).

4. IRENA and Methanol Institute. Innovation Outlook: Renewable Methanol. (International Renewable Energy Agency, 2021)

5. Liu, Z., Wang, K., Chen, Y., Tan, T. & Nielsen, J. Third-generation biorefineries as the means to produce fuels and chemicals from CO_2_. Nat. Catal. 3, 274−288 (2020).

6. Guo, Y. et al. Methylotrophy of *Pichia pastoris*: Current Advances, Applications, and Future Perspectives for Methanol-Based Biomanufacturing. ACS Sustainable Chem. Eng. 10, 1741–1752 (2022).

7. Guo, F., Qiao, Y., Xin, F., Zhang, W. & Jiang, M. Bioconversion of C1 feedstocks for chemical production using *Pichia pastoris*. Trends Biotechnol. 41, 1066−1079 (2023).

8. Bhataya, A., Schmidt-Dannert, C. & Lee, P.C. Metabolic engineering of *Pichia pastoris* X-33 for lycopene production. Process Biochem. 44, 1095−1102 (2009).

9. Liu, Y. et al. Engineered monoculture and co-culture of methylotrophic yeast for de novo production of monacolin J and lovastatin from methanol. Metab. Eng. 45, 189−199 (2018).

10. Gao, J., Li, Y., Yu, W. & Zhou, Y.J. Rescuing yeast from cell death enables overproduction of fatty acids from sole methanol. Nat. Metab. 4, 932−943 (2022).

11. Cai, P. et al. Methanol biotransformation toward high-level production of fatty acid derivatives by engineering the industrial yeast *Pichia pastoris*. Proc. Natl. Acad. Sci. U. S. A. 119, e2201711119 (2022).

12. Cos, O., Ramón, R., Montesinos, J.L. & Valero, F. Operational strategies, monitoring and control of heterologous protein production in the methylotrophic yeast *Pichia pastoris* under different promoters: a review. Microb. Cell Fact. 5, 17 (2006).

13. Werpy, T. & Petersen, G. Top Value Added Chemicals from Biomass: Volume I--Results of Screening for Potential Candidates from Sugars and Synthesis Gas. (United States, 2004).

14. Ji, H. et al. Preparation of bio-based elastomer and its nanocomposites based on dimethyl itaconate with versatile properties. Composites Part B: Engineering 248, 110383 (2023).

15. Guo, B. et al. Biobased Poly(propylene sebacate) as Shape Memory Polymer with Tunable Switching Temperature for Potential Biomedical Applications. Biomacromolecules 12, 1312−1321 (2011).

16. Wang, R., Zhang, J., Kang, H. & Zhang, L. Design, preparation and properties of bio-based elastomer composites aiming at engineering applications. Compos. Sci. Technol. 133, 136−156 (2016).

17. Wang, Y., Guo, Y., Cao, W. & Liu, H. Synergistic effects on itaconic acid production in engineered *Aspergillus niger* expressing the two distinct biosynthesis clusters from *Aspergillus terreus* and *Ustilago maydis*. Microb. Cell Fact. 21, 158 (2022).

18. Elkasaby, T., Hanh, D.D., Kawaguchi, H., Kondo, A. & Ogino, C. Effect of different metabolic pathways on itaconic acid production in engineered *Corynebacterium glutamicum*. J. Biosci. Bioeng. 136, 109−116 (2023).

19. Feng, J. et al. Construction of cell factory through combinatorial metabolic engineering for efficient production of itaconic acid. Microb. Cell Fact. 21, 275 (2022).

20. Rong, L. et al. Engineering *Yarrowia lipolytica* to Produce Itaconic Acid From Waste Cooking Oil. Front. Bioeng. Biotechnol. 10, 888869 (2022).

21. Zhao, C. et al. Enhanced itaconic acid production in *Yarrowia lipolytica* via heterologous expression of a mitochondrial transporter MTT. Appl. Microbiol. Biotechnol. 103, 2181−2192 (2019).

22. Cai, P. et al. Recombination machinery engineering facilitates metabolic engineering of the industrial yeast *Pichia pastoris*. Nucleic Acids Res. 49, 7791−7805 (2021).

23. Palmieri, L. et al. Identification of the Yeast Mitochondrial Transporter for Oxaloacetate and Sulfate. J. Biol. Chem. 274, 22184−22190 (1999).

24. Farré, J.-C., Li, P. & Subramani, S. BiFC Method Based on Intraorganellar Protein Crowding Detects Oleate-Dependent Peroxisomal Targeting of *Pichia pastoris* Malate Dehydrogenase. Int. J. Mol. Sci. 22(2021).

25. Luo, Z. et al. Enhanced pyruvate production in *Candida glabrata* by carrier engineering. Biotechnol. Bioeng. 115, 473−482 (2018).

26. Tang, W.-Y. et al. Metabolic engineering of *Yarrowia lipolytica* for improving squalene production. Bioresour. Technol. 323, 124652 (2021).

27. Koivistoinen, O.M. et al. Glycolic acid production in the engineered yeasts *Saccharomyces cerevisiae* and *Kluyveromyces lactis*. Microb. Cell Fact. 12, 82 (2013).

28. Berrios, J. et al. Role of Dissimilative Pathway of *Komagataella phaffii* (*Pichia pastoris*): Formaldehyde Toxicity and Energy Metabolism. Microorganisms 10, 1466 (2022).

29. Yu, Y.-f., Yang, J., Zhao, F., Lin, Y. & Han, S. Comparative transcriptome and metabolome analyses reveal the methanol dissimilation pathway of *Pichia pastoris*. BMC Genomics 23, 366 (2022).

30. Wu, X. et al. Efficient Bioproduction of 3-Hydroxypropionic Acid from Methanol by a Synthetic Yeast Cell Factory. ACS Sustainable Chem. Eng. 11, 6445–6453 (2023).

31. Lin-Cereghino, G.P. et al. Mxr1p, a key regulator of the methanol utilization pathway and peroxisomal genes in *Pichia pastoris*. Mol. Cell Biol. 26, 883–97 (2006).

32. Wang, X. et al. Mit1 Transcription Factor Mediates Methanol Signaling and Regulates the Alcohol Oxidase 1 (AOX1) Promoter in *Pichia pastoris*. J. Biol. Chem. 291, 6245–61 (2016).

33. Jastroch, M., Divakaruni, A.S., Mookerjee, S., Treberg, J.R. & Brand, M.D. Mitochondrial proton and electron leaks. Essays Biochem. 47, 53−67 (2010).

34. Xi, Y. et al. Metabolic engineering of the acid-tolerant yeast *Pichia kudriavzevii* for efficient L-malic acid production at low pH. Metab. Eng. 75, 170−180 (2023).

35. Canales, C., Altamirano, C. & Berrios, J. Effect of dilution rate and methanol-glycerol mixed feeding on heterologous *Rhizopus oryzae* lipase production with *Pichia pastoris* Mut^+^ phenotype in continuous culture. Biotechnol. Prog. 31, 707−714 (2015).

36. Lim, C.K. et al. Designing and Engineering *Methylorubrum extorquens* AM1 for Itaconic Acid Production. Front. Microbiol. 10, 1027 (2019).

37. Sonntag, F., Buchhaupt, M. & Schrader, J. Thioesterases for ethylmalonyl–CoA pathway derived dicarboxylic acid production in *Methylobacterium extorquens* AM1. Appl. Microbiol. Biotechnol. 98, 4533−4544 (2014).

38. Quynh Le, H.T., Anh Mai, D.H., Na, J.-G. & Lee, E.Y. Development of *Methylorubrum extorquens* AM1 as a promising platform strain for enhanced violacein production from co-utilization of methanol and acetate. Metab. Eng. 72, 150−160 (2022).

39. Zhu, W.-L. et al. Bioconversion of methanol to value-added mevalonate by engineered *Methylobacterium extorquens* AM1 containing an optimized mevalonate pathway. Appl. Microbiol. Biotechnol. 100, 2171−2182 (2016).

40. Sonntag, F. et al. Engineering *Methylobacterium extorquens* for de novo synthesis of the sesquiterpenoid α-humulene from methanol. Metab. Eng. 32, 82−94 (2015).

41. Yuan, X.-J. et al. Rewiring the native methanol assimilation metabolism by incorporating the heterologous ribulose monophosphate cycle into Methylorubrum extorquens. Metab. Eng. 64, 95–110 (2021).

42. Naerdal, I., Pfeifenschneider, J., Brautaset, T. & Wendisch, V.F. Methanol-based cadaverine production by genetically engineered *Bacillus methanolicus* strains. Microb. Biotechnol. 8, 342–50 (2015).

43. Drejer, E.B. et al. Methanol-based acetoin production by genetically engineered *Bacillus methanolicus*. Green Chem. 22, 788−802 (2020).

44. Chowdhury, N.P., Litty, D. & Müller, V. Biosynthesis of butyrate from methanol and carbon monoxide by recombinant *Acetobacterium woodii*. Int. Microbiol. 25, 551−560 (2022).

45. Litty, D. & Müller, V. Butyrate production in the acetogen *Eubacterium limosum* is dependent on the carbon and energy source. Microb. Biotechnol. 14, 2686−2692 (2021).

46. Humphreys, J.R. et al. Establishing *Butyribacterium methylotrophicum* as a Platform Organism for the Production of Biocommodities from Liquid C_1_ Metabolites. Appl. Environ. Microbiol. 88, e0239321 (2022).

47. Yu, W., Gao, J., Yao, L. & Zhou, Y.J. Bioconversion of methanol to 3-hydroxypropionate by engineering *Ogataea polymorpha*. Chin. J. Catal. 46, 84−90 (2023).

48. Li, J. et al. Engineering yeast for high-level production of β-farnesene from sole methanol. Metab. Eng. 85, 194−200 (2024).

49. Bachleitner, S., Severinsen, M.M., Lutz, G. & Mattanovich, D. Overexpression of the transcriptional activators *Mxr1* and *Mit1* enhances lactic acid production on methanol in *Komagataella phaffii*. Metab. Eng. 85, 133−144 (2024).

50. Luo, G. et al. Overproduction of Patchoulol in Metabolically Engineered *Komagataella phaffii*. J. Agric. Food. Chem. 71, 2049−2058 (2023).

